# A protocol for cortical and deep-brain calcium imaging in freely moving mice with the Mini2P system

**DOI:** 10.64898/2026.11.09.764980

**Authors:** Irma A. Tello-García, Sammy Alhassen, Carlos Portera-Cailliau, William Zeiger

## Abstract

Two-photon calcium imaging has transformed our understanding of neuronal circuit function, yet conventional approaches require head fixation, limiting the study of neural activity during natural behavior. Miniaturized two-photon microscopy (Mini2P) overcomes this constraint by enabling calcium imaging in freely moving animals while preserving the optical single-cell resolution of conventional two-photon microscopy. However, its implementation requires the careful integration of multiple technically demanding procedures, presenting substantial challenges for laboratories adopting this approach. Here, we present a comprehensive, step-by-step protocol for implementing Mini2P calcium imaging in freely moving mice, emphasizing procedures that are unique to or particularly critical for successful implementation of this technique. This protocol covers system preparation and performance checks, optical alignment, field-of-view selection, baseplate installation, and strategies for maintaining imaging stability during microscope attachment and reattachment. We also address behavioral habituation, integration with behavioral tasks, imaging acquisition, quality control, and troubleshooting as a part of an integrated experimental workflow. Representative imaging data acquired through cranial windows in the primary visual and somatosensory cortices, and through a gradient-index lens in the medial prefrontal cortex, illustrate the applicability of this workflow across different brain regions and optical preparations. Designed for adaptation across brain regions and behavioral paradigms, the protocol provides practical guidance for identifying technical challenges and verifying successful implementation. By establishing clear decision points, validation criteria, and troubleshooting strategies at each stage, this workflow aims to improve reproducibility and facilitate the adoption of the Mini2P for investigating neuronal circuit dynamics during freely moving behavior.

## INTRODUCTION

Two-photon (2P) calcium imaging (2PCI) is a powerful technique for recording neuronal activity in vivo in rodents and other species. In vivo 2PCI has traditionally required animal head-fixation and the use of a large stationary benchtop microscope, a constraint incompatible with naturalistic, self-paced behaviors such as exploration or social interactions (Grienberger et al., 2022). This limitation motivated years of efforts to engineer lightweight miniaturized head-mounted microscopes for 2PCI in freely behaving rodents (Flusberg et al., 2005; Zong et al., 2017, 2021, 2022; Ozbay et al., 2018; Madruga et al., 2025). Early designs were not broadly adopted because of issues with fiber dispersion, slow scanning, excessive weight, and small fields of view (FOV). More recent implementations, especially the Mini2P design (Zong et al., 2022), have overcome several of these constraints: it is ultra-low-weight (<2.4 g), has a highly flexible and durable fiber-optic cable, supports stable, multiplane imaging of neurons, and, using successive stitching of adjacent FOVs allows recordings from more than 10,000 neurons in the same animal, all without impeding freely-moving behavior (Zong et al., 2022). Moreover, three interchangeable objectives extend the use of the Mini2P to different preparations from standard glass-covered cranial windows to more invasive prisms and gradient-index (GRIN) lenses, enabling deep brain imaging (Zong et al., 2022).

The Mini2P was released as a fully open-source platform along with extensive supplemental information that described the optical and mechanical design of the microscope, with component lists, mechanical drawings, 3D models, assembly instructions, video tutorials, acquisition and analysis software, imaging parameters, and performance tests, a credit to the developers (Zong et al., 2022; Kavli-NTNU, n.d.). Day-to-day operational knowledge is available through an open-access repository (https://github.com/kavli-ntnu/MINI2P_toolbox), videos describing the microscope assembly and calibration, and in-person workshops hosted by the developing laboratory (KavliInstituteforSystemsNeuroscience, n.d.). However, although these amazing resources are publicly available and openly shared, the information can be daunting for researchers less familiar with 2P microscopy or instrumentation. Thus, many laboratories may find this exciting technology challenging to implement. Moreover, hands-on training is currently only available to those attending limited capacity workshops in Norway. This gap has grown even more apparent as integrated turnkey Mini2P systems have become commercially available through multiple vendors (Inscopix, LABmaker, PhenoSys, ThorLabs). To our knowledge, these commercial options do not come with an operational protocol that matches the specific hardware that a purchasing laboratory receives. In short, the current lack of detailed step-by-step instructions may limit the widespread adoption of the Mini2P technology.

Here, we address this gap with a detailed step-by-step protocol for implementing Mini2P calcium imaging in freely moving mice, with an emphasis on the experimental steps that are particularly critical for the Mini2P. Our detailed instructions cover the following steps: alignment of the microscope to the optical plane, identification and documentation of the selected FOV, positioning and cementing of the baseplate, verification of FOV stability after cementing, microscope attachment and re-attachment, animal habituation, cable management, freely moving acquisition, quality control, and troubleshooting in a single end-to-end protocol. Because they are not covered here in any detail, we invite the reader to refer to other protocols for general instructions on how to carry out survival surgeries for cranial window implantation, viral delivery, and GRIN lens implantation, or on how to train rodents on behavioral assays, or analyze calcium imaging data (Mostany and Portera-Cailliau, 2008; Barretto et al., 2009; Holtmaat et al., 2009; Yang et al., 2010; Low et al., 2014; Resendez et al., 2016; Zhang et al., 2019; Grienberger et al., 2022; Zhao et al., 2025). Those procedures should be selected and validated for the target brain region, cell population and scientific question.

The presented workflow is intended to support laboratories either extending established in vivo imaging workflows or adopting the Mini2P for the first time. Potential applications include longitudinal recordings of large neuronal populations during spatial navigation, sensory-guided behavior, learning, memory retrieval, decision-making, social interactions and motor behaviors, as well as comparison of circuit dynamics across experimental manipulations and disease models. This protocol is designed to be broadly applicable across brain regions and behavioral paradigms rather than tied to a single cortical area or task. By defining explicit decision points, success criteria, and troubleshooting actions at each stage, the workflow aims to improve reproducibility while preserving the high resolution, large-scale neuronal recordings that the Mini2P enables during self-directed behavior.

## METHODS

### 1. Experimental animals

All experiments followed the U.S. National Institutes of Health guidelines for animal research, under animal use protocols (ARC#2007-035 and ARC#2020-119) approved by the Chancellor’s Animal Research Committee and Office for Animal Research Oversight at the University of California, Los Angeles. All experiments used male and female C57BL/6J mice, 5–8 weeks old at the time of surgery. We crossed the TIGRE2-jGCaMP8s-IRES-tTA2-WPRE (JAX line 037952) and Slc17a7-IRES2-Cre-D line (JAX 037512) to generate heterozygous double transgenic animals with stable GCaMP8s expression in excitatory neurons. All mice were housed in a vivarium with food and water provided *ad libitum* (except during behavior training, see below) and an inverted 12/12 h light/dark cycle (experiments were performed during the dark cycle).

### 2. Custom 3D-printed components

Some components of our Mini2P experimental setup are 3D-printed and were designed or modified in our laboratory to facilitate day-to-day experiments (**Fig. 1A-C**).

**Figure 1.**
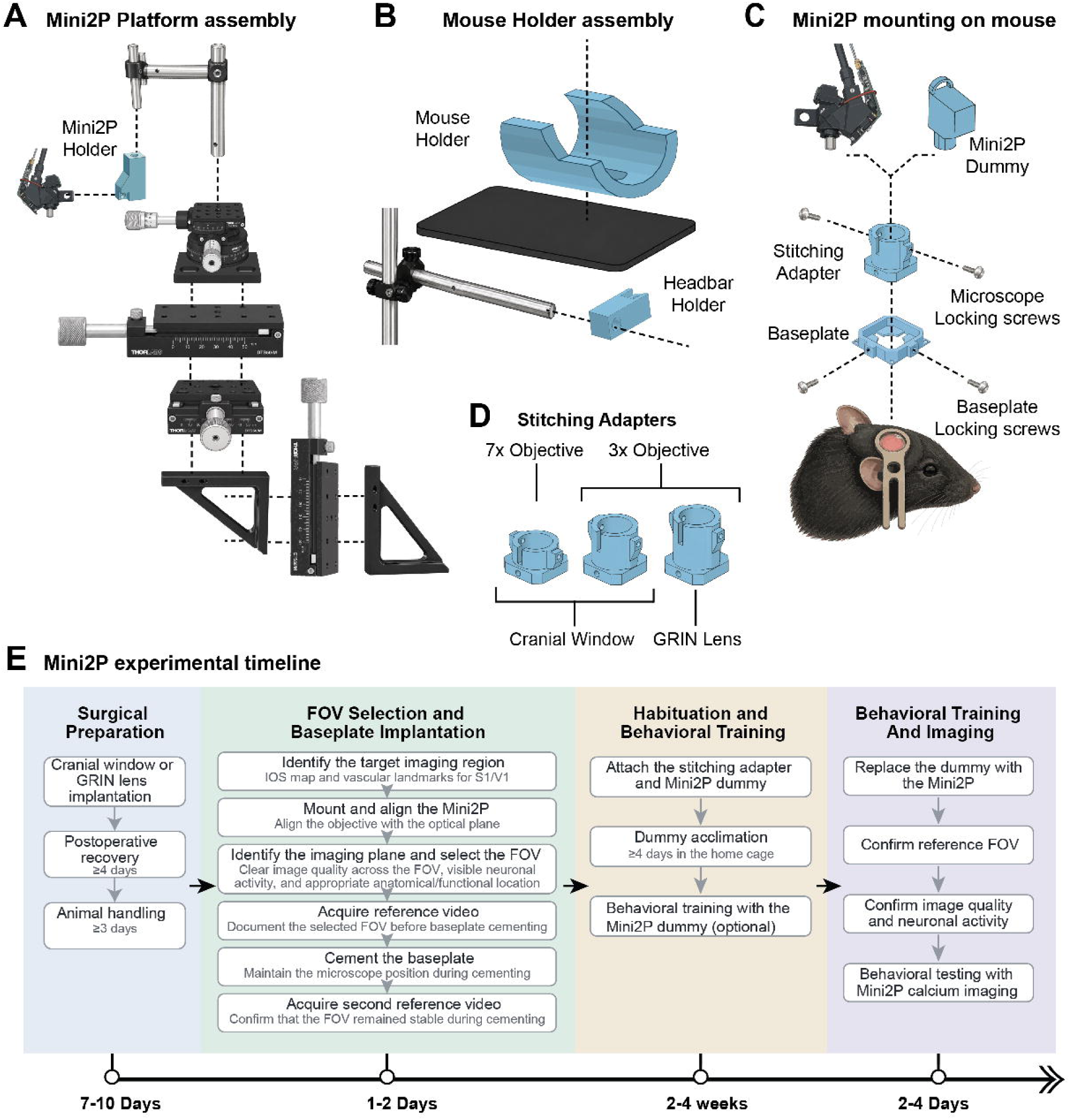
Mini2P mounting hardware and experimental workflow. **(A)** Expanded schematic of the Mini2P mounting platform, consisting of three translation stages for independent adjustment along the x-, y-, and z-axes, a goniometric stage for angular adjustment, and a custom-designed Mini2P holder used to support and position the microscope during mounting. Components colored in blue in panels A-D are 3D printed. **(B)** Assembly of the custom-designed mouse holder and headbar holder. The mouse is positioned within the holder, and the implanted headbar is secured to the headbar holder using a locking screw to stabilize the animal during Mini2P alignment and FOV selection. **(C)** Expanded schematic illustrating the assembly of the Mini2P microscope, stitching adapter, and baseplate onto the implanted headbar. The stitching adapter is secured to the baseplate using two baseplate locking screws that engage with corresponding threaded holes in the stitching adapter, while a separate set of microscope locking screws secures the Mini2P or dummy within the stitching adapter. **(D)** Three modified stitching adapters designed for different objectives and imaging preparations, including the 7x and 3x objectives for imaging through cranial windows and the 3x objective for imaging through an implanted GRIN lens. **(E)** Schematic timeline illustrating the major stages of a Mini2P experiment, including surgical preparation, FOV selection and baseplate implantation, habituation and behavioral training, and behavioral testing with calcium imaging. Approximate durations are indicated for each stage and may vary depending on the surgical preparation and behavioral paradigm.

#### 2.1. Stitching Adapter

In most cases, the stitching adapter remains attached to the animal between imaging sessions since it is also used to hold the Mini2P dummy during habituation and behavioral experiments. We made several modifications to the original stitching adapter design. The baseplate itself follows the unmodified design described by Zong et. al. (2022). To improve reproducibility, we added two threaded holes at the bottom of the stitching adapter that align with the existing baseplate locking screws. Rather than pressing against the adapter, these screws now thread directly into the new holes, allowing them to be fully tightened without displacing or deforming the stitching adapter (**Fig. 1C**). This secures the adapter in the same position within the baseplate across imaging sessions. We also incorporated fixed screw stops for the microscope locking screws that secure the Mini2P body within the stitching adapter (**Fig. 1C**). The fixed stops allow the microscope locking screws to be fully tightened without relying on the experimenter to judge the screw depth, resulting in more consistent microscope positioning. We also changed the orientation of the microscope locking screws. In the original design, the screws contact the microscope body at an angle, which does not provide consistent contact with the cylindrical surface of the scope. Our redesigned adapter moves the screws to opposite sides of the microscope, producing a more direct and symmetric clamping force (**Fig. 1C**). Finally, we designed separate stitching adapters for the 3x and 7x Mini2P objectives. The original design positions both objectives at the same height despite their different working distances. Because the 7x objective has a much shorter working distance, we redesigned its adapter to position the objective closer to the cranial window. The height of the 3x adapter was also reduced, as the original design positioned the objective slightly too far from the window surface. These objective-specific changes provide a more appropriate starting position and greater working range for each objective (**Fig. 1D**).

#### 2.2. Headbar holder

The headbar holder was designed in-house to mount into a standard optical post, secured by an 8-32 screw. This holder secures the headbar implanted on the mouse head and is used to head-fix the mouse during Mini2P mounting, repositioning, and removal (**Fig. 1B**).

#### 2.3. Mini2P holder

The Mini2P holder was also designed in-house to support the Mini2P when it is not mounted on the head of the mouse and to position the Mini2P during the mounting procedure, providing stable positioning during FOV selection (**Fig. 1A**).

#### 2.4. Mouse holder

This piece was designed as an open cylindrical shape that attaches to a flat base with a screw. The animal is positioned inside the mouse holder during head fixation to minimize body movements and provide additional stability during the Mini2P mounting procedure without interfering with access to the implanted headbar and cranial window (**Fig. 1B**).

#### 2.5. Mini2P dummy

We made several modifications to the original Mini2P dummy design. This 3D-printed version allows us to have several Mini2P dummies available for multiple experimental mice. Our version has a hollow interior and a removable lid. Small weights are placed inside to achieve a final weight of 3 g after which the lid is permanently attached. We additionally incorporated a loop into the Mini2P dummy from which a lightweight cord can be attached, allowing the mice to be habituated not only to the head-mounted weight but also to the presence and movement of a tether approximating the Mini2P optic fiber during behavioral training (**Fig. 1C**).

## STEP-BY-STEP PROTOCOL

We developed a modular mounting platform using commercially available positioning stages and custom-designed components to facilitate Mini2P mounting and FOV selection. The platform incorporates three translation stages (x-y-z adjustments) and a goniometric stage for angular adjustment of the microscope (**Fig. 1A**). A custom-designed mouse holder and headbar holder further allow the Mini2P to be positioned over the cranial window or implanted GRIN lens (**Fig. 1A**). The Mini2P is secured to the animal through a baseplate and stitching adapter assembly, which also allows the microscope to be removed and reattached between imaging sessions (**Fig. 1A**). We modified the stitching adapter to improve the consistency of microscope positioning and developed separate adapters for 3x and 7x objectives and GRIN lens preparations (**Fig. 1D**).

The overall experimental workflow is summarized in **Fig. 1E**. Although the duration of each stage may vary depending on the imaging preparation and behavioral task, the same general sequence can be followed across cortical and deep-brain imaging experiments. Throughout the detailed protocol below we provide recommendations for common challenges; comprehensive troubleshooting can be found in **Table 2**.

### 1. Surgical Preparation

#### 1.1. Cranial Window Surgery

- For Mini2P calcium imaging in primary somatosensory cortex (S1) or primary visual cortex (V1), the first step is to surgically implant a cranial window (∼4 mm diameter coverslip) over the target cortical region in anesthetized mice. Detailed protocols for cranial window surgery have been previously published (Mostany and Portera-Cailliau, 2008; Holtmaat et al., 2009). [Note: personal preferences regarding coverslip shape and size (or stacking) for the window should not affect Mini2P imaging, although this was not assessed here; we used single round #1 size coverslips, 4 mm in diameter]
- Monitor anesthesia depth repeatedly throughout the procedure by observing respiratory rate and pinch reflex responses.
- Always administer Carprofen (5 mg/kg, i.p., Zoetis) and dexamethasone (0.2 mg/kg, i.p., Vet One) before surgery for analgesia and to reduce brain edema, respectively.

#### 1.2. GRIN lens implantation for deep-brain imaging

- For deep-brain Mini2P imaging, for example in the medial prefrontal cortex (mPFC) optical access requires implantation of a gradient-refractive-index (GRIN) relay lens rather than a standard cranial window. The detailed surgical procedure for GRIN lens implantation has been described previously (Zhao et al., 2025).
- Following skull preparation, a craniotomy matching the diameter of the GRIN lens (1 mm in our case) is created over the target coordinates. The overlying tissue can then either be aspirated to the desired implantation depth, as described (Zhao et al., 2025), or a blunt needle can be slowly lowered to the target depth to create an insertion track without tissue aspiration. When using the latter approach, the needle is withdrawn and the GRIN relay lens is subsequently lowered along the preformed track to the desired depth. Following either approach, the lens is secured with cyanoacrylate adhesive, followed by headbar implantation and reinforcement with dental acrylic.

#### 1.3. Prism implantation for lateral optical access

- Mini2P imaging is also compatible with implanted microprisms, which can provide optical access to brain regions in fissures or tissue planes that are difficult to image through a conventional cranial window. Although prism-based imaging was not performed in the experiments described here, established protocols for microprism implantation (Low et al., 2014) can be adapted for use with the Mini2P system.

#### 1.4. Headbar implantation

- After coverslip sealing, glue a custom-made headbar (modified from M2P Headbar-0d, LabMaker) to the skull surrounding the coverslip using cyanoacrylate adhesive (**Fig. 2A**). Although animals are freely moving during imaging, the headbar is used to stabilize the mice during intrinsic signal imaging, Mini2P alignment, field-of-view (FOV) selection, baseplate cementing and Mini2P mounting for behavioral sessions.
- Take care to ensure correct positioning of the headbar (**Fig. 2A**).
- Once the adhesive is dry, apply black dental cement (Lang Dental Mfg. Co., Inc.) around the headbar and the edges of the coverslip (**Fig. 2A**). Using black dental cement helps reduce light contamination during imaging. Avoid applying excess dental cement over the headbar.

#### 1.5. Recovery and post-operative monitoring

- After surgery, allow mice to recover on a heating pad until fully ambulatory (30-60 min).
- Administer Carprofen (5 mg/kg, i.p., Zoetis) post-operatively every 24 hours for 3 days.
- Monitor all animals daily for signs of pain or distress, abnormal locomotion, poor grooming, weight loss, wound disruption, implant instability, or window opacity.
- Allow mice to recover for at least 4 days before beginning habituation to handling, the behavioral apparatus, and Mini2P mounting.

### 2. Intrinsic Signal Imaging

To identify S1 and V1 prior to 2PCI consider using intrinsic signal imaging (ISI) to guide FOV selection for the Mini2P. The detailed procedures have been published previously (Zeiger et al., 2021; Vasquez et al., 2023). For ISI in S1, we mapped the location of the barrel field, 5-7 days after cranial window surgery, as described previously (Dobler et al., 2024). Mice were head-fixed in a custom-made 3D-printed headbar holder (Bamboo Lab X1 Carbon printer; **Fig. 1B**). All contralateral whiskers were gently attached with bone wax to a glass capillary tube coupled to a ceramic piezoelectric bending actuator (PI127-Physik Instrumente) (**Fig. 2B, left**). To map the location of V1, visual stimulation was provided by a single flashing white LED in front of the contralateral eye. Responses for 30 stimulation trials were averaged; each consisted of a 5 Hz LED flashing stimulus for 1.5 s. The response signal, divided by the averaged baseline signal and summed across all trials, was used to generate the visual cortical map as described previously (Vasquez et al., 2023) (**Fig. 2B, right**).

**Figure 2.**
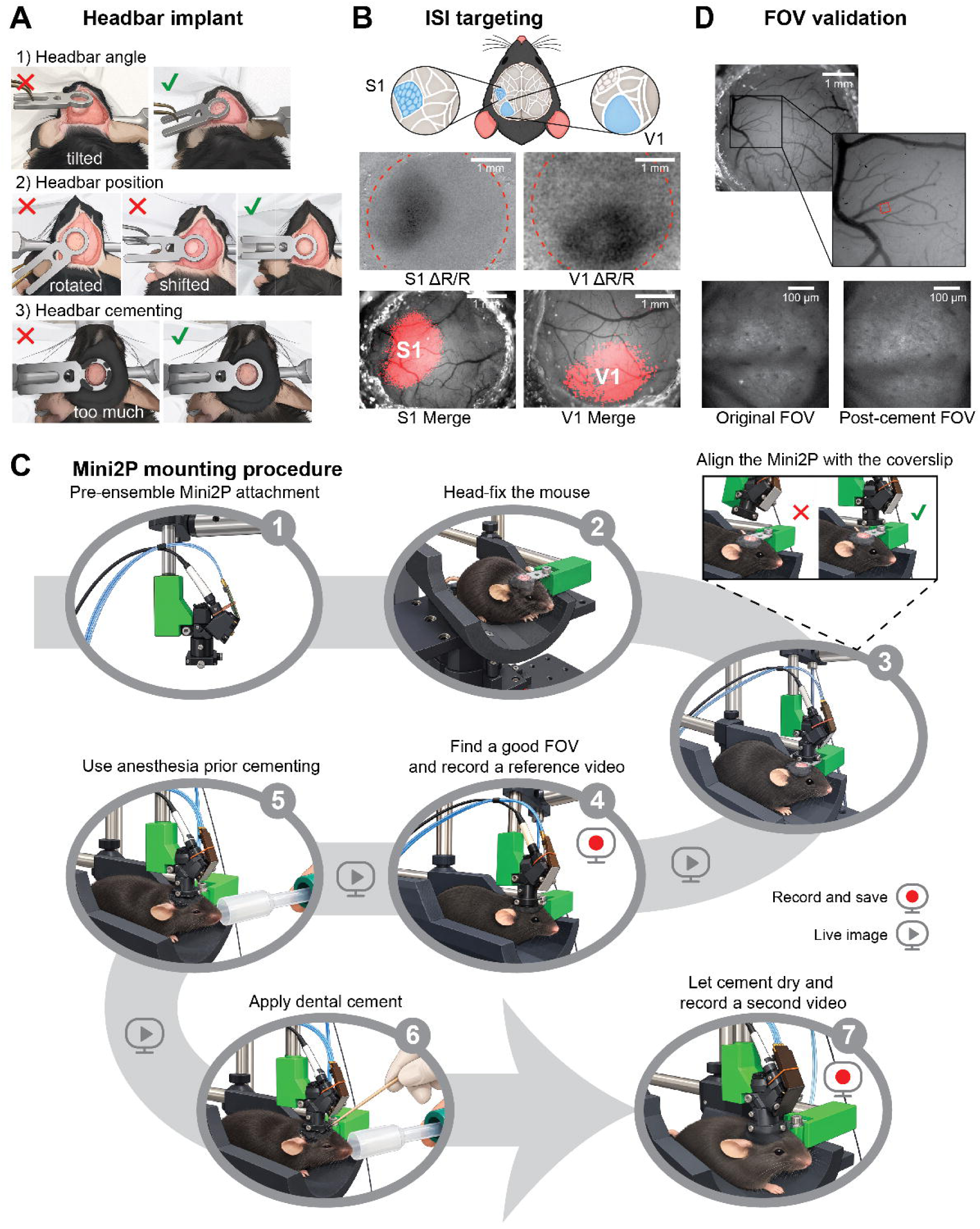
Surgical preparation, FOV targeting, and Mini2P mounting procedure. **(A)** Examples of correct and incorrect headbar implantation. (1) The headbar should be positioned parallel to the plane of the cranial window to facilitate subsequent Mini2P alignment. (2) The headbar should be oriented approximately perpendicular to the mouse’s body axis and centered around the cranial window. (3) Avoid excessive dental cement extending above the headbar, which can interfere with baseplate positioning. **(B)** Intrinsic signal imaging (ISI) used to functionally identify the S1 and V1 cortices. Representative ISI response maps (ΔR/R) and their corresponding overlays with cortical vasculature are shown. These maps are subsequently used to guide Mini2P positioning and FOV selection within the desired cortical region. **(C)** Step-by-step Mini2P mounting procedure. (1) The Mini2P attachment (Mini2P body + stitching adapter + baseplate) is assembled before mounting on the animal’s head. (2) Position the animal in the mouse holder and secure the implanted headbar. (3) Align the Mini2P objective parallel to the cranial window. (4) Identify a suitable FOV using live imaging and acquire a reference recording. (5) Maintain the animal under light anesthesia during preparation for cementing. (6) Apply dental cement around the baseplate while maintaining microscope position. (7) After the cement cures and the animal recovers from anesthesia, acquire a second reference recording to verify FOV stability. **(D)** FOV identification and validation during Mini2P mounting. Cortical vascular landmarks are used to guide microscope positioning. Example calcium imaging fields acquired before and after baseplate cementing show corresponding anatomical structures, illustrating maintenance of the selected FOV through the mounting procedure.

For mPFC experiments, ISI imaging was not performed. Instead, the final imaging field of view was selected during Mini2P mounting as described below.

### 3. Mini2P mounting and Field-of-View (FOV) selection

Mounting of the Mini2P microscope onto the head of the animal is achieved via the baseplate and stitching adapter. The baseplate serves as a structure to securely position and mount the Mini2P body and stitching adapter to the skull. The stitching adapter couples the Mini2P to the baseplate, allowing the microscope position and angle to be adjusted even after the baseplate is permanently fixed to the mouse head (Zong et al., 2022). Together, these components ensure consistent microscope positioning and reproducible imaging of the same FOV across sessions. Once a suitable FOV is identified, the baseplate is cemented onto the implanted headbar and remains in place for the duration of the experiment

3.1. The Mini2P, stitching adapter and baseplate should be assembled before mounting on the animal’s head (**Fig. 1C and 2C, step 1**). The modified stitching adapter fits snugly inside the baseplate and is secured with two baseplate locking screws that thread directly into the corresponding holes at the bottom of the adapter. Fully tighten both screws to secure the stitching adapter in a consistent position. The Mini2P then fits into the stitching adapter and is secured using a second set of microscope locking screws with fixed screw stops. This allows the microscope to be repeatedly removed and remounted while maintaining the selected FOV.
3.2. Anesthetize the animal (we use isoflurane 3% induction, 1.5% maintenance) prior to head fixation to a custom-built platform using the implanted headbar and a custom-made and 3D-printed headbar holder (**Fig. 2C, step 2**;). We use a platform that consists of three 50-mm dovetail translation stages (Thorlabs, DTS50/M), which provide independent adjustment along the x-, y-, and z-axes and a rotation goniometer pair (Thorlabs, GNL20/M) for angular adjustment, assembled using a metric stage angle bracket (Thorlabs, DTSA03/M). Detailed instructions for assembling the mounting stage are provided in **Figure 1A**.
3.3. Align the cranial window and Mini2P microscope in the same plane (**Fig. 2C, step 3**). First, lower the Mini2P along the z-axis as close as possible to the cranial window and headbar without allowing the microscope to contact either structure. Next, use the x-axis for lateral adjustment and the y-axis for anterior–posterior adjustment to center the Mini2P objective over the center of the coverslip. Once centered, adjust the tilt of the Mini2P with the goniometer until the objective is parallel to the plane of the coverslip. This alignment is critical for obtaining a uniform FOV and must be based on the angle of the coverslip rather than that of the headbar. Finally, make small adjustments along all axes as necessary to refine the alignment while maintaining sufficient clearance between the microscope and the cranial window.
3.4. Once the objective is positioned over the target region, initiate live imaging. Mini2P image acquisition is controlled using the software ScanImage (**Fig. 2C, step 4**). Manually lower the Mini2P using the z-axis of the mounting platform. This should be performed slowly and in very small increments while continuously monitoring live imaging to avoid contact between the objective and the cranial window. Once the surface of the cortex appears on live imaging, advance using even smaller increments until neuronal somata become visible. For imaging in S1 or V1, use the ISI map and vascular landmarks visible through the cranial window to identify the cortical region of interest and align the Mini2P objective with the center of the functionally identified region.
3.5. Adjust the imaging plane in the x-and y-axes to identify a suitable FOV. Use the software-controlled z-axis in ScanImage for fine adjustments in small increments. Select a FOV based on uniform image intensity throughout the FOV, clearly resolved neuronal somata, and detectable spontaneous or evoked neuronal activity during live imaging. To evoke neural activity while looking for a FOV in S1, we use a cotton swab to stimulate the contralateral whiskers during live imaging (**Fig. 2C, step 4**).
3.6. After the desired FOV and imaging depth are identified, stop live imaging and acquire a short reference video before fixing the Mini2P. This video is used later to compare with a post-cementing recording and is helpful to detect any FOV displacement caused by mouse movement or accidental Mini2P displacement during the cementing procedure (**Fig. 2C, step 4**).
3.7. Cement the Mini2P baseplate to the skull using black dental cement while maintaining the microscope position. During this step, keep mice under light isoflurane anesthesia (1.5-2%) to minimize movements that are often triggered by the scent of the dental cement (**Fig. 2C, step 5**).
3.8. Apply the cement carefully around the baseplate to provide a stable attachment, taking extreme care not to obstruct the optical path or contact the cranial window (**Fig. 2C, step 6**).
  - Using a thin wooden applicator stick, collect a small drop of dental cement and place it onto the edge of the baseplate.
  - With the same stick, carefully spread the cement toward the existing headcap surrounding the cranial window.
  - Repeat this process around the entire baseplate, adding one drop at a time until the dental cement covers the whole surface and there are no areas where light can leak inside. During this process, it is important to keep the dental cement out of the coverslip and away from the eyes and whiskers of the mouse. Ensure that the cement contacts only the baseplate and does not touch the stitching adapter or the Mini2P body, as this could bond the microscope assembly to the baseplate and prevent its subsequent removal.
3.9. After the cement has fully cured, which usually takes approximately 15 minutes, discontinue anesthesia.
3.10. Once mice are fully awake, acquire a second reference video and compare it with the pre-cement recording to confirm that the FOV has not shifted during baseplate cementing (**Fig. 2C, step 7; Suppl. Fig. 1**). A slight displacement can be corrected by adjusting the microscope locking screws that secure the Mini2P body to the stitching adapter while doing live imaging to identify the desired FOV. However, large shifts caused by mouse movement or accidental displacement of the Mini2P are generally irreversible once the cement is fully cured. If substantial movement occurs during fixation, do not complete the cementing process; instead, reposition the microscope using live imaging and repeat the FOV-selection procedure described in paragraphs 3.4-3.6 above.
3.11. After FOV stability is confirmed, remove the Mini2P microscope and replace it with a custom-made 3D-printed Mini2P dummy microscope (3 g) that is subsequently used to habituate the mouse to the weight of the Mini2P.

### 4. Habituation and animal handling

General handling, baseplating, and acclimation to the Mini2P dummy were performed similarly across cortical targets and deep-brain imaging experiments.

4.1. General handling and Mini2P dummy acclimation

- Following postoperative recovery, handle mice with gloved hands for at least 5 minutes per day for a minimum of 3 consecutive days to reduce stress associated with handling.
- Following handling, head-fix mice for Mini2P mounting, FOV selection, and baseplate implantation as described above.
- After baseplate implantation, secure the stitching adapter to the baseplate and then attach the 3D-printed Mini2P dummy. The fully weighted dummy is left in place continuously for at least 4 days to acclimate animals to the additional head-mounted weight before imaging.
- Continuously monitor the animals during this period and proceed to behavioral training and imaging once they exhibit normal movement and behavior while carrying the dummy.
4.2. S1 behavioral training

- For the imaging experiments described here, mice were habituated to the behavioral arena for 10 minutes on 2 consecutive days. The arena consisted of a linear-track (50 x10 cm, corridor width 5 cm) built from 3D-printed components, with a lickport at each end and four textured panels in the middle of the track.
- Water-restricted mice were trained to travel back and forth between the two lickports, receiving water rewards after licking at a rewarded lickport illuminated with an LED.
- Animals were trained until they successfully obtained rewards on >80% of trials. Training was then continued while animals carried the Mini2P dummy to allow them to adjust their movement and head orientation to the additional weight
- Imaging sessions were initiated once animals moved normally through the arena while carrying the dummy and again achieved >80% task accuracy, which typically required 2–3 additional training sessions.

### 5. Image acquisition

The following steps describe how calcium imaging data is collected using the Mini2P microscope during freely moving behavior. We used ScanImage Basic 2023.1.0 for microscope acquisition (Pologruto et al., 2003).

5.1. Before each recording session, attach the Mini2P to the implanted baseplate using the stitching adapter. Identify once again the previously selected field of view (FOV) by comparing live imaging with the reference images acquired during baseplate implantation. As described in Section 3.10, small shifts can be corrected by adjusting the screws that secure the Mini2P body to the stitching adapter. If the Mini2P shifts slightly during reattachment, loosen both screws and reposition the Mini2P under live imaging until the desired FOV is recovered. Then, tighten the two microscope locking screws to their fixed stops, alternating between them while continuously monitoring live imaging. Do not force the screws beyond their fixed stops or apply so much force as to displace the Mini2P. Continue until the microscope is securely attached and the desired FOV remains stable. An example of successful re-attachment across sessions is shown on **Suppl. Fig. 1**.
5.2. Confirm image quality and neuronal activity before beginning behavioral testing. Acquire recordings at 14.6 Hz, with laser power typically ranging from 65-80 mW, as measured through the microscope. Adjust laser power as needed to obtain clear neuronal signals while avoiding saturation. Acquire images at 256 x 256 pixels. Although we use the parameters described above, framerate and image acquisition size can be adjusted based on experimental needs.
5.3. For cortical imaging through cranial windows in S1 and V1, select the desired imaging plane within the superficial cortex based on calcium indicator expression (e.g., 120-200 μm below the dura for layer 2/3), image quality, density and activity of labeled neurons, and the anatomical or functional location of the FOV.
5.4. For deep-brain imaging through an implanted GRIN lens, focus the imaging plane at the GRIN lens surface.
5.5. Acquisition procedures are otherwise similar across the 3x, 7x, and GRIN preparations, although the 7x objective provides a substantially smaller FOV (500 x 500 μm for 3x and 250 x 250 μm for 7x).
5.6. Determine the recording duration based on the animal’s progression through the behavioral task.

### 6. System synchronization

To synchronize Mini2P calcium imaging, behavioral events, and video camera data streams (controlled by the open-source behavior acquisition hardware Bpod through TTL signals), we used the data acquisition hardware integrated with the Mini2P system. A Vidrio vDAQ connected the Mini2P acquisition system with the Bpod controller and the externally triggered behavioral camera. The camera was connected by USB to a separate Dell laptop used for behavioral tasks control and camera acquisition (**Table 1**). Bpod OUT1 was connected to vDAQ D1.0, configured for ScanImage acquisition start and stop on rising edges. vDAQ D0.1 was connected to both Bpod OUT2 and the camera through a shared connection, while vDAQ D3.7 was connected to Bpod IN1. (**Suppl. Fig. 2A**). These connections allowed TTL signals to coordinate the imaging and behavioral acquisition systems.

6.1. To initiate synchronized recordings, configure the behavioral camera to start upon receiving an external trigger (**Suppl. Fig. 2B**). In ScanImage, enable external acquisition start/stop triggering and select LOOP to await the trigger (**Suppl. Fig. 2C**). Initiate the MATLAB/Bpod behavioral script, which sends the trigger to begin Mini2P acquisition and behavioral video recording. These signals provide common reference events for temporally aligning calcium imaging with behavior.
6.2. During behavioral testing, predefined task events detected by Bpod (or other behavioral task control hardware), such as licking for a reward at the lickport, to generate a trigger that stops the current recording/trial and immediately initiates a new trial/recording. In our linear-track task, only a correct lick at the rewarded port or time out will trigger the next trial and initiate the stop/start transition.
6.3. Mini2P acquisition was therefore saved as a sequence of separate trial recordings, rather than as a single continuous imaging movie. This helps to keep imaging file sizes more manageable, but continuous imaging throughout a task is possible as well.
6.4. The start and stop events, together with Bpod event timestamps, provide common temporal reference points that can be used subsequently to align Mini2P imaging, behavioral video, and behavioral events across the session.

**Table 1.** Materials/Equipment.

| Equipment/ Components | Source/ Vendor | Catalog no./file |
|---|---|---|
| **Animals** |  |  |
| B6.slc17a7 x jgcamp8 | The Jackson Laboratory | JAX stock #037512 × JAX stock #037952 |
| **Mini2P Imaging System** |  |  |
| Mini2P and Control Box v1 | Thorlabs | FP: 16296 336776 |
| 3x objective | Thorlabs | D3X-5W1 |
| 3x objective (GRIN) | Thorlabs | D3X-5A1 |
| 3x objective (Prism) | Thorlabs | D3X-45P1 |
| 7x objective | Thorlabs | D7X-7W1 |
| Laser | Spark Lasers | ALCOR 920-Mini2P |
| **Mounting Hardware** |  |  |
| Large Dual-Axis Goniometer | Thorlabs | GNL20/M |
| Dovetail Stage (x3) | Thorlabs | DTS50/M |
| Angle Bracket | Thorlabs | DTSA03/M |
| Dummy weights (316 Stainless Steel Washer for M6 Screw) | McMaster-Carr | 90965A170 |
| Headbar | SendCutSend | Mini2P_HeadbarLong_SA.dxf |
| Baseplate & stitching adapter screws | McMaster-Carr | 92015A205; 91067A111 |
| Headbar screw (Socket Head Cap, #5-44 x 3/8", SS 316) | Fastenere | SHCS316SS544 |
| **Custom 3D-Printed Components** |  |  |
| Bambu Lab X1-C 3D Printer | Bambu Lab | PF001-PSA001-U |
| PLA Matte Black Filament | Bambu Lab | 11101 |
| Mini2P holder | In-house (custom) | Mini2P_Holder.stl |
| Stitching adapter (3x) | In-house (modified) | Mini2P_3x_StitchingAdapter.stl |
| Stitching adapter (7x) | In-house (modified) | Mini2P_7x_StitchingAdapter.stl |
| Stitching adapter (7x GRIN) | In-house (modified) | Mini2P_7x_StitchingAdapter_GRIN.stl |
| Dummy | In-house (custom) | Mini2P_Dummy.stl |
| Headbar holder | In-house (custom) | Mini2P_HeadbarHolder.stl |
| Mouse holder assembly | In-house (custom) | Mini2P_MouseHolder.stl |
| **Behavioral and Acquisition Equipment** |  |  |
| Bpod State Machine r2+ | Sanworks | 1036 |
| FLIR Blackfly S | Teledyne Vision Solutions | BFS-U3-23S3M-C |
| Mini2P acquisition computer | Supplied with Mini2P system (Thorlabs) | Intel Core i5-12600k (12th Gen); 32GB RAM; NVIDIA GeForce GTX 1660 SUPER (6GB); Windows 11 |
| Behavioral acquisition laptop | Dell | Latitude 5540; 13th Gen Intel Core i7-1370P; 32 GB RAM; Windows 11 Enterprise |
| Data acquisition hardware | Supplied with Mini2P system (Thorlabs) | – |
| **Software** |  |  |
| Image/Fiji | Open source | v1.54 |
| AutoCAD | Autodesk | v24.2 |
| Suite2p (Python) | Open source | v1.1.1 |
| Spinnaker | Teledyne Vision Solutions | Version x.x |
| MATLAB | MathWorks | R2023b |
| Spark Lasers GUI | Spark Lasers | v19.1.9 |
| ScanImage (MATLAB) | MBF Bioscience | v2023.1.0 |

### 7. Basic analysis pipeline

Raw Mini2P movies were first visually inspected for image quality, motion artifacts, and stability of the imaging FOV.

7.1. Calcium imaging data processing
  - Calcium imaging data were processed using Suite2p for motion correction and automated segmentation of neuronal regions of interest (ROIs).
  - Identified ROIs were subsequently manually curated to remove non-neuronal or poorly segmented ROIs. Data is then exported to MATLAB for subsequent analysis.
  - Neuropil-corrected fluorescence traces were calculated as:

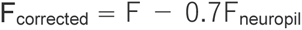
  - For subsequent analysis, fluorescence traces were normalized using a modified z-score. For each ROI, a moving window was used to identify the period of lowest fluorescence variability. The mean and standard deviation within this minimum-variance window were used as the baseline mean and standard deviation, respectively, and the complete fluorescence trace was normalized relative to this baseline. Since the length of the trials may vary significantly, the baseline mean was set as 1/10 of the trial length.
7.2. Behavioral data processing
  - Behavioral data recorded by Bpod were processed to extract behavioral events, and their corresponding timestamps.
  - Behavioral videos were analyzed using DeepLabCut (DLC;(Mathis et al., 2018) to track the nose, neck, left and right ears, center of the head, two points along the body, and the base of the tail.

DLC-derived tracking data and Bpod behavioral events were subsequently aligned with the calcium imaging data, allowing neuronal activity to be analyzed in relation to task events and animal movement.

## RESULTS

### FOV targeting and validation during Mini2P mounting

Successful Mini2P imaging requires appropriate positioning of the headbar and microscope to ensure optical access and stable mounting. We identified several aspects of headbar implantation that are important for subsequent Mini2P mounting (**Fig. 2A**), including positioning the headbar parallel to the cranial window (**Fig. 2A 1**), maintaining a perpendicular orientation relative to the mouse’s body axis (**Fig. 2A, step 2**), and avoiding excess dental cement that could interfere with baseplate positioning(**Fig. 2A, step 3**). These preparations facilitate alignment of the Mini2P objective with the cranial window and subsequent FOV selection.

For cortical imaging, ISI was used to functionally identify S1 and V1 before Mini2P mounting (**Fig. 2B**). Functional maps were overlaid onto images of cortical vasculature to provide anatomical landmarks for locating the target cortical region during subsequent FOV selection.

The mounting procedure is summarized in **Fig. 2C**, beginning with microscope and animal positioning, followed by optical alignment, FOV selection, and baseplate cementing. Particular attention was given to aligning the microscope with the coverslip plane to achieve uniform focus across the FOV before baseplate fixation. A second reference recording was acquired after cement curing and recovery from anesthesia. Representative pre- and post-cement recordings show that the same imaging field and anatomical landmarks can be maintained during baseplate fixation (**Fig. 2D**). An example of the stability of a selected FOV across two subsequent imaging sessions following microscope removal and reattachment can be found in **Suppl. Fig. 1**. These observations provide qualitative examples of repeatable FOV recovery, without establishing longitudinal identification of the same individual neurons.

### Mini2P calcium imaging across different brain regions and optical preparations

To demonstrate the application of the Mini2P across different brain regions and optical configurations, we acquired calcium imaging recordings from freely moving mice in S1, V1, and mPFC (**Fig. 3**). Representative maximum-intensity projections illustrate the imaging FOVs obtained using the 3x objective in S1 (**Fig. 3A**). Individual neuronal ROIs were identified using Suite2p (**Fig. 3B**). We then extracted and z-scored fluorescence traces from neuronal ROIs and identified clearly detectable calcium transients with excellent signal-to-noise ratios during freely moving behavior (**Fig. 3C-D**). Similar results were obtained in V1 with the 3x objective (**Fig. 3E-H**). Higher resolution imaging was obtained using the 7x objective in S1 (**Fig. 3I-L**). Deep-brain imaging in mPFC was also achieved using the 7x objective and an implanted GRIN lens (**Fig. 3M-P**). Together, these examples demonstrate that the Mini2P can be used to obtain cellular-resolution calcium imaging recordings across different cortical regions, optical configurations, and deep-brain preparations using the experimental workflow described above.

**Figure 3.**
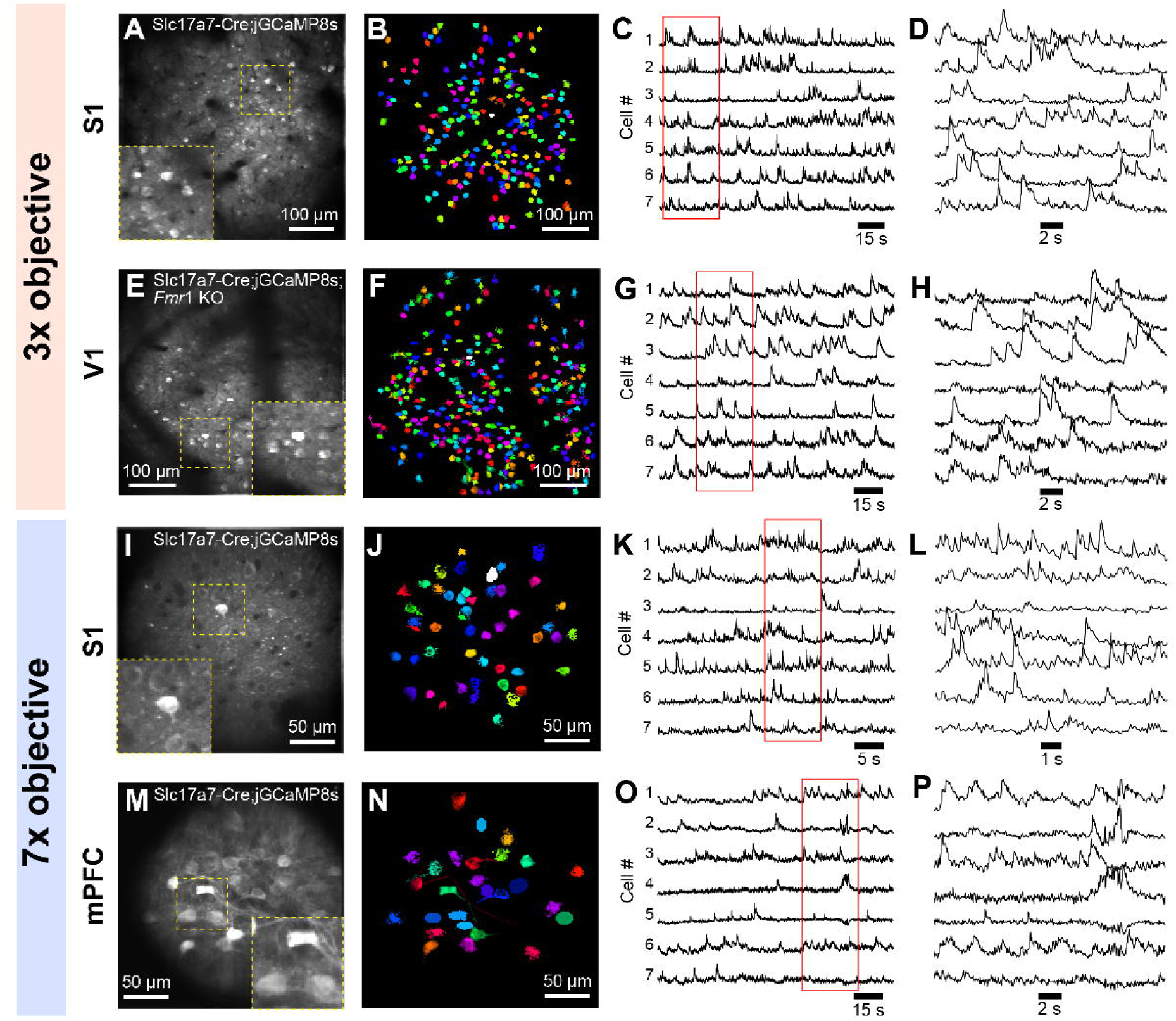
Representative Mini2P calcium imaging recordings across cortical and deep-brain preparations. **(A–D)** Representative calcium imaging of GCaMP8s-expressing neurons in S1 using the 3x objective. (A) Maximum-intensity projection of the recorded imaging FOV, with an enlarged view of the region indicated by the yellow dashed box. (B) Neuronal soma ROIs identified using Suite2p and subsequent manual curation. (C) Representative calcium activity traces from seven example neurons, with the red box indicating the time interval expanded in (D). **(E–H)** Calcium imaging in V1 using the 3x objective. Panels show the maximum-intensity projection (E), identified neuronal ROIs (F), representative calcium traces (G), and an expanded recording interval (H). **(I–L)** Higher-magnification calcium imaging in S1 using the 7x objective. Panels show the maximum-intensity projection (I), identified neuronal ROIs (J), representative calcium traces (K), and an expanded recording interval (L). **(M–P)** Deep-brain calcium imaging in mPFC through an implanted GRIN lens using the 7x objective. Panels show the maximum-intensity projection (M), identified neuronal ROIs (N), representative calcium traces (O), and an expanded recording interval (P). All recordings were acquired in freely moving mice. Calcium activity traces are shown as modified z-scored fluorescence signals. Each trace was normalized to its peak value.

### Simultaneous calcium imaging and behavioral tracking during freely moving behavior

To demonstrate the integration of Mini2P calcium imaging with behavioral measurements, we acquired synchronized recordings of neuronal activity and animal movement during a behavioral task performed along a linear corridor (**Fig. 4**). Animal position was tracked using DLC and aligned with the corresponding calcium imaging recordings. Representative examples illustrate the position of the animal and neuronal activity within the same FOV during periods of low activity, high activity, and drinking (**Fig. 4A**). Individual neuronal ROIs were color-coded according to their z-scored fluorescence activity, allowing visualization of changes in neuronal activity at each selected behavioral time point.

**Figure 4.**
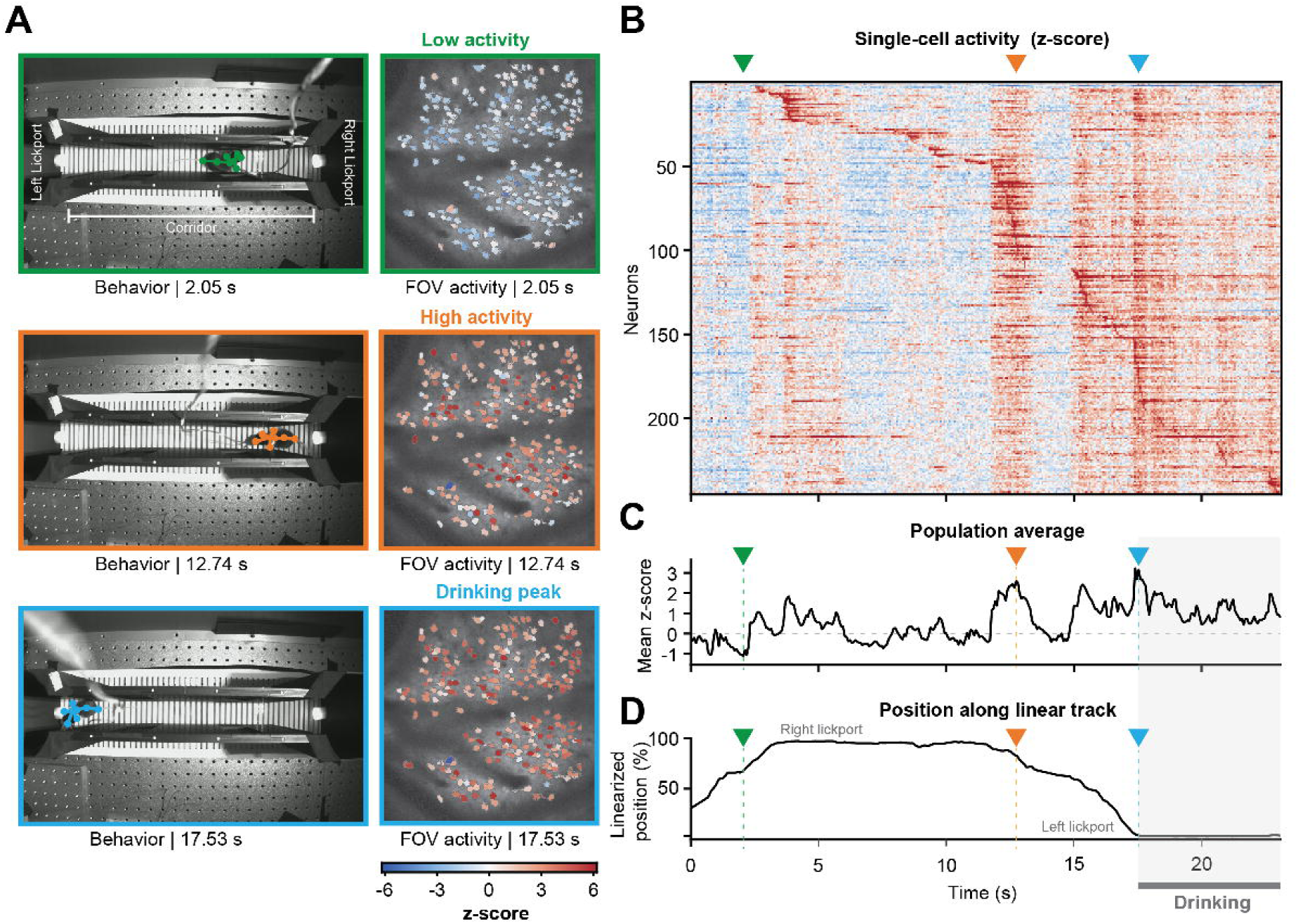
Simultaneous Mini2P calcium imaging and behavioral tracking in freely moving mice. **(A)** Representative examples of synchronized behavioral tracking and neuronal calcium activity during a period of low neuronal activity (green, 2.05 s), high neuronal activity (orange, 12.74 s), and drinking (blue, 17.53 s). Behavioral images show the position of the animal along a linear corridor, tracked using DLC, with lickports located at either end. Corresponding Mini2P imaging FOVs show segmented neuronal ROIs, color-coded according to their z-scored fluorescence activity at each selected time point. **(B)** Heatmap of z-scored fluorescence activity from individual neurons throughout the recording. Each row represents a neuron, and color indicates the relative fluorescence activity over time. **(C)** Mean z-scored fluorescence activity across the recorded neuronal population over the same interval. **(D)** Linearized position of the animal along the behavioral corridor over time, with 0% and 100% corresponding to the left and right lickports, respectively. Colored arrows in (B–D) indicate the corresponding low-activity (green), high-activity (orange), and drinking (blue) time points illustrated in (A). Gray horizontal bars indicate the drinking period.

The corresponding calcium imaging data were used to examine individual neuronal activity and the mean activity across the recorded population over time (**Fig. 4B,C**). These recordings were aligned with the animal’s linearized position along the corridor, allowing changes in neuronal activity to be examined in relation to movement between the two lickports and periods of drinking (**Fig. 4D**). Together, these examples demonstrate the ability to synchronize Mini2P calcium imaging with behavioral tracking and relate neuronal activity to specific behavioral events during freely moving experiments.

## DISCUSSION

The relatively recent improvements in the design of miniaturized head-mounted microscopes for 2PCI in freely behaving rodents (Zong et al., 2017, 2021, 2022; Ozbay et al., 2018; Madruga et al., 2025) are a long-awaited development in neuroscience. When coupled with naturalistic behavior assays, this technology will help uncover more ethologically relevant circuit dynamics. The publication of the original Mini2P system provided extensive open-source information for microscope construction and operation (Zong et al., 2022). However, the Mini2P is not quite “plug- and-play,” and successful laboratory implementation depends on a sequence of experimental and technical procedures to ensure stable, reproducible imaging. Here, we address this gap by providing a protocol that incorporates a detailed sequence of experimental and technical procedures into a practical and reproducible experimental workflow for new Mini2P users. Beyond providing step-by-step instructions, our protocol emphasizes decision points, confirmation criteria, and corrective strategies at stages where errors can still be reversible. The intention is to make the practical knowledge necessary for successful Mini2P implementation more explicit and transferable, reducing reliance on trial-and-error learning or direct training from laboratories already experienced with the technique.

A key feature of this workflow is its generalizability across brain regions. Although surgical access and optical implants must be tailored to the specific target structure, the core experimental sequence for imaging remains largely unchanged across preparations. Accordingly, the examples of cortical imaging and deep-brain imaging presented here illustrate how the same general workflow can be used while adapting the surgical and optical components to access a particular brain structure. Although the present protocol was developed and validated using the commercially available Thorlabs Mini2P, which was based directly on the original Mini2P design described by Zong et. al. (2022) and is therefore applicable to laboratories building their own systems, most of the steps are not limited to this specific microscope architecture and should generalize to other systems. Establishing reproducible alignment between the microscope and the imaging surface, selecting and documenting the FOV, minimizing mechanical displacement during mounting and remounting, habituating animals to the weight of the Mini2P, synchronizing imaging with behavioral videos, and verifying FOV stability before data acquisition are general requirements for reliable Mini2P calcium imaging. These steps of the workflow could therefore be transferred to other similar commercial systems; however, specific hardware, such as the stitching adapter, Mini2P holder, and headbar holder, may require small adjustments to be compatible with other available systems.

An additional strength of the approach described here is its modular, DIY implementation. Rather than requiring laboratories to reproduce a single preset hardware configuration or purchase all peripheral components as a complete system, several elements of our setup were designed or modified in-house and then printed using a 3D printer, assembled from commercially available components, or machined by independent companies. These include the Mini2P and headbar holder, the Mini2P dummy, headbar, baseplates, and stitching adapters, as well as components of the behavioral arena and synchronization setup. Beyond the cost-saving benefits, this strategy allows laboratories to tailor the experimental setups to their available space, existing equipment, and behavioral requirements while preserving the mechanical stability and optical alignment required for reliable Mini2P imaging. Local fabrication and modular assembly make it possible to modify individual components as experimental requirements evolve in an easy and fast way, integrate Mini2P with existing laboratory equipment, and replace or redesign peripheral hardware without restructuring the entire imaging system. Thus, the implementation described here provides a flexible framework in which commercial, off-the-shelf, and custom-built components can be combined according to the needs and resources of each laboratory.

A further contribution of our protocol is the systematic troubleshooting framework (**Table 2**) developed around common failure points in Mini2P experiments. Many problems encountered during imaging originate from steps prior to image acquisition and can therefore be difficult to diagnose without explicit checkpoints. Our troubleshooting framework organizes each potential problem and its likely cause, and then provides a corrective or preventive action, as well as a verification criterion indicating whether the problem has been resolved. This structure is intended to help users distinguish potentially reversible implementation errors from those that will require removing an experimental animal from experiments, thus reducing time spent on troubleshooting.

**Table 2.** Troubleshooting for Mini2P Experiments. Potential problems that may arise during Mini2P experiments and recommended corrective actions.

| Experimental phase | Problem | Possible cause | Solution | Verification |
|---|---|---|---|---|
| Headbar Implantation | The plane of the headbar is not parallel to the coverslip. This makes the positioning of the Mini2P challenging or ultimately makes it impossible to image. | The headbar was positioned tilted relative to the plane of the window during headbar implantation (**Fig. 2Aa**). | Before cement is fully cured, readjust the headbar to ensure it is positioned completely parallel to the coverslip plane. | The headbar and coverslip planes are parallel, and the headbar remains in this position after cement cures (**Fig. 2Aa**). |
| Headbar Implantation | The headbar is rotated relative to the mouse’s body. This could make head-fixation difficult | The headbar was positioned rotated towards the torso of the mouse (**Fig. 2Ab**). | Before cementing, rotate the headbar until it is perpendicular (90º) to the mouse’s body axis. Reposition the headbar before dental cement cures. | The headbar is perpendicular to the mouse’s body axis and can be easily head-fixed to the platform (**Fig. 2Ab**). |
| Headbar Implantation | The headbar partially covers the coverslip, partially obstructing the cranial window and limiting access for Mini2P mounting. | The headbar was positioned off-center relative to the coverslip (**Fig. 2Ab**). | Before applying dental cement, reposition the headbar so that the coverslip is centered within the headbar opening. If misalignment is detected during cementing, correct the headbar position before the dental cement cures. | The coverslip is centered within the headbar opening, and the cranial window remains fully visible for Mini2P mounting (**Fig. 2Ab**). |
| Headbar Implantation | The Mini2P baseplate cannot be fully lowered onto the headbar. | Excess dental cement was applied, and it extends above the upper surface of the headbar, obstructing the baseplate and preventing it from being placed above the headbar (**Fig. 2Ac**). | Before the dental cement cures, remove the excess cement and reshape the headcap so that the cement remains below the upper surface of the headbar and does not obstruct the baseplate contact area. If the cement is already cured, do not force the baseplate into position; remove the obstructing cement before proceeding with Mini2P mounting. | The upper surface of the headbar is unobstructed, and the Mini2P baseplate can be fully lowered into its intended position without contacting the dental cement (**Fig. 2Ac**). |
| Recovery | The mouse shows persistent pain, weight loss, poor grooming, or abnormal locomotion. | Surgical complications, such as infection or inability to tolerate the implant/Mini2P. | Do not begin habituation or imaging. Evaluate the animal according to the approved postoperative monitoring protocol. Remove mice from further experiments if they don’t recover properly. | Proceed only after the animal has recovered and moves normally. |
| Mini2P mounting | The cranial window becomes opaque, bloody or blood vessels and tissue are no longer clearly visible. | Tissue growth, bleeding, debris, or deterioration of the window. | Re-evaluate the window before mounting the Mini2P. Exclude mice from imaging if window quality is not good. | Only use mice when the window is clear and free from bone growth and blood vessels are visible. |
| Mini2P mounting | The objective cannot be centered over the target region. | Insufficient travel along the x- or y-axis, incorrect initial positioning of the mouse, or interference from the headbar or headcap. | Retract the Mini2P along the z-axis, reposition the mouse on the mounting platform making sure the coverslip is parallel to the Mini2P axis. Repeat x- and y-axis alignment before lowering the Mini2P. | The objective is centered over the target area with sufficient clearance from the headbar and cranial window. The Mini2P axis is parallel to the coverslip. |
| Mini2P mounting | The Mini2P objective contacts the cranial window, headbar, or dental cement. | Clearance was not checked from multiple angles, or the objective was tilted incorrectly and the Mini2P was lowered too quickly. | Retract the Mini2P. Check the position of the mouse, re-position lateral and angular alignment before approaching the window again in smaller z-axis increments while continuously monitoring live imaging. If the problem is not solved after this, please check the next row below. | The cortical surface appears in live imaging without physical contact between the microscope and the implant. |
| Objective and adapter selection | The focal plane (e.g. cortical surface) cannot be reached within the available physical range. | Incorrect stitching adapter for the objective or inappropriate initial objective-to-window distance (**Table 3**). | Confirm that the stitching adapter corresponds to the 3x, 7x objective or GRIN lens being used. Make changes if needed (**Table 3**). | The cortical surface can be brought into focus without the objective being in contact with the cranial window or GRIN lens. |
| Objective and adapter selection with GRIN lenses | Mini2P objective is too far from the GRIN lens to achieve focus or contacts the lens directly during positioning | GRIN lens length and implantation depth affect the distance between the lens surface and Mini2P objective. The provided stitching adapter is optimized for our specific GRIN lens and mPFC preparation and may not be ideal for other configurations | Adjust the stitching adapter height to accommodate the GRIN lens length and implantation depth for the intended imaging preparation. | Confirm that the objective can be positioned at the appropriate working distance without contacting the GRIN lens and that a clear FOV can be obtained. |
| FOV selection | Only part of the FOV is in focus or image intensity is uneven across the FOV. | The objective is not parallel to the coverslip. | Retract slightly and adjust the goniometer to align the objective with the plane of the coverslip rather than the plane of the headbar. Repeat fine x-, y-, and z-axis adjustments as needed. | Live image is focused across the whole FOV. |
| FOV selection | Brain tissue is visible, but neuronal bodies cannot be found. | The imaging plane is too superficial or deep; the calcium indicator was not expressed properly or was injected outside the imaging depth. | Use small physical z-axis adjustments until tissue is visible, then use the software-controlled z-axis in ScanImage for fine focusing. | Individual neuronal somata are clearly visible and remain in focus throughout the FOV. |
| FOV selection | The image is visible, but spontaneous neuronal activity cannot be detected. Cells look filled, but they don't show activity. | The imaging plane contains few labeled neurons; stimulation is required to detect activity; AAVs expression is too high. | Explore nearby imaging planes with more labeled neurons. If neurons are not spontaneously active, stimulate according to the brain region (i.e., deflect whiskers for S1) to confirm neuronal activity in the area. Do not select the FOV based solely on anatomical appearance. If AAVs expression is too high, remove the mouse from further experiments. | Neuronal somata exhibit detectable spontaneous or evoked activity during live imaging. |
| Baseplate cementing | The animal moves while dental cement is being applied. | Aversion to the scent of the dental cement or insufficient anesthesia. | Stop cement application before fixation is completed. Confirm anesthesia at 1.5-2%, recover the FOV under live imaging, repeat the FOV-selection and take another FOV reference video procedure. | The FOV matches the pre-cement reference and remains stable while cement is applied. |
| Baseplate cementing | Dental cement spreads toward the cranial window, eyes, or whiskers. | Excess cement was applied, the mixture was too fluid, or the cement was deposited too close to these structures. | Stop adding cement and prevent further spread by removing dental cement while it is not cured. Apply only small drops to the baseplate edge and spread each drop toward the existing headcap. | The cranial window and optical path remain unobstructed, and no cement contacts the eyes or whiskers. |
| Baseplate cementing | Cement contacts the stitching adapter or Mini2P body, preventing Mini2P removal from the mouse head. | Cement was spread beyond the edge of the baseplate. | Stop cement application and remove uncured excess taking care of not moving the Mini2P. Do not force removal of the Mini2P if cement has already been cured around it. | After curing, the stitching adapter and Mini2P can be removed independently from the permanently fixed baseplate. |
| Baseplate cementing | Light enters around the baseplate during imaging. | There are unfilled gaps between the baseplate and the headbar. | After confirming that the FOV remains stable, add small drops of black dental cement to the uncovered areas. | No visible light contamination is present in the live image. |
| Baseplate cementing | A small FOV shift is observed between the pre- and post-cement reference videos. | Minor displacement of the Mini2P body within the stitching adapter. | Loosen the screws securing the Mini2P body to the stitching adapter and reposition the microscope under live imaging. Retighten the two screws gradually, alternating between them one turn at a time. | The original FOV is recovered and remains stable after both screws are secured. |
| Baseplate cementing | A big FOV shift occurs during cementing. A large shift is not recoverable by adjusting the stitching adapter screws after the cement has cured. | Substantial animal movement during baseplate cementing. | If the cement is not fully cured, stop cementing, reposition the microscope under live imaging, and repeat FOV selection. | The recovered FOV matches the pre-cement reference, or a new FOV is selected before cementing resumes. |
| Mini2P Dummy habituation | The mouse shows abnormal posture, reduced movement, or tries to remove the dummy constantly. | Insufficient habituation. | This is normal during the first days of habituation. Continue the habituation and delay imaging until normal movement and head posture are restored. | The animal moves, grooms, and explores normally while carrying the dummy in a natural head position. |
| Mini2P reattachment | The FOV differs between the reference video and Mini2P reattachment. | Asymmetric tightening of the stitching adapter’s screws, or a change in the Mini2P position. | Under live imaging, loosen the stitching adapter’s screws, recover the reference FOV and retighten them gradually, alternating between them until the Mini2P is securely attached and the FOV matches the reference video. | Imaging FOV matches the reference video recorded. |
| Mini2P reattachment | Tightening the screws displaces the FOV or deforms the stitching adapter. | Overtightening of stitching adapter screws | Loosen the screws and reposition the Mini2P under live imaging. Tighten gradually and symmetrically, stopping once the microscope is secure. Avoid additional tightening after fixation is achieved. | Gentle handling of Mini2P does not move the microscope, and the FOV remains unchanged. |
| Behavioral Training | Task performance worsens after attachment of the Mini2P dummy. | The mouse has not adapted its movement and head position to the added weight. | Continue behavioral training with the dummy in place until the performance threshold is achieved. | The mouse moves normally and is able to reach the set performance threshold while carrying the dummy. |
| Behavior and Mini2P recordings | The optic fiber becomes twisted, tangled, or restricts the mouse’s movement. | Cable rotation accumulated during the session, inappropriate cable support or counterweighting. | Stop the experiment, remove the mouse from the arena, untwist and reposition the cable, return the mouse, and resume only after unrestricted movement is restored. | The animal can reach all relevant areas of the arena without visible cable tension or movement restriction. |
| Behavior and Mini2P recordings | The mouse moves less than during training sessions or avoids part of the arena. | Cable tension due to insufficient suspended length. | Stop the experiment and inspect cable suspension and available length; confirm sufficient cable length and restart the experiment. | Mouse can move normally across all the arena areas. |
| Behavior and Mini2P recordings | The FOV moves abruptly during acquisition. | Mechanical displacement of the Mini2P due to loose screws. | Stop the session and manually inspect the Mini2P for movement. If present, head-fix the mouse and, under live imaging, tighten the screws to completely secure the Mini2P in place. Confirm manually that it is not moving anymore before restarting recordings. | The FOV remains stable during gentle manual movement of Mini2P. |
| Behavior and Mini2P recordings | Brief white flashes appear across the FOV during recordings when the mouse is moving. | Light leakage: external light is reaching the objective through gaps around the baseplate, headcap or stitching adapter. | Stop the recording, check the screws securing the Mini2P body to the stitching adapter and the screws locking the stitching adapter to the baseplate. If necessary, tighten them. Inspect the baseplate-headcap junction and seal any visible gaps with small amounts of black dental cement. Allow the cement to cure before restarting the recordings. | No white flashes or light changes are observed in the FOV while recording. |

**Table 3.** Suggested setups for different imaging applications.

| Imaging Application | Objective | NA | Working Distance | Stitching Adapter | Application/Considerations |
|---|---|---|---|---|---|
| Large FOV/cortical imaging | 3x | 0.5 | 1.17mm | 3x adapter | Large FOV population imaging through a cranial window (example of S1 in **Fig. 3A-D** and V1 in **Fig. 3E-H**). |
| Higher magnification/subcellular imaging | 7x | 0.7 | 0.41mm | 7x adapter | Smaller FOV population imaging through a cranial window (example of S1 in **Fig. 3I-L**). Higher magnification and better spatial resolution could be used for subcellular imaging (e.g., dendrites, axons), but shorter working distance complicates positioning and finding suitable FOV. |
| GRIN lens | 7x | 0.7 | 0.41mm | 7x adapter (GRIN) | Deep brain population imaging through an implanted GRIN lens (example of mPFC imaging in **Fig. 3M-P**). |
| Prism | 3x | 0.5 | 2.32mm | 3x adapter | Population imaging across cortical layers through an implanted prism. |

Despite these advantages, Mini2P remains a technically challenging approach. Reliable recordings depend on high-quality optical implants, careful alignment, stable fixation of the mounting components, and appropriate animal habituation. Maintaining the same FOV across longitudinal sessions can also be difficult because small changes in microscope position can alter the imaging plane. Other factors, such as microscope weight, cable tension, and attachment stability, are relevant during freely moving recordings since they can affect animal movement or introduce imaging artifacts. In addition, although downstream analysis is not addressed in detail here, integrating neuronal activity with behavioral videos and synchronizing task events represents an important challenge that should be considered when designing the experimental and analysis pipeline.

Taken together, the protocol presented here provides more than a description of how to operate a particular Mini2P system. It provides a practical framework for implementing Mini2P imaging across different brain regions while preserving the core steps required for stable and reproducible recordings. By combining modular hardware, explicit validation criteria, and troubleshooting strategies, this workflow aims to make the technical knowledge that is usually acquired through trial and error more accessible to new users. This may lower practical barriers to adopting the Mini2P as an imaging technique while allowing laboratories to adapt the system to their own biological questions, behavioral paradigms, and available infrastructure.

## Acknowledgements

This work was supported by the National Institute of Child Health and Development (R01HD108370 and R01HD054453 to C.P.-C.), National Institute of Neurological Disorders and Stroke (1R01NS129517-01 to W.Z.), the Marion Bowen Neurobiology Postdoctoral Scholar Grant Program (to I.A.T.-G.), the American Parkinson’s Disease Association (1268719 to WZ and 1452632 to SA) and the UCLA David Binder Fund for Lewy Body Disease Research. During the preparation of this manuscript, the authors used ChatGPT generative AI image tools (OpenAI; GPT-5.6 Sol) to create cartoon illustrations based on original laboratory photographs, as well as to assist with grammar on the early drafts of the manuscript. All AI-assisted content was reviewed and revised by the authors to ensure scientific accuracy and faithful representation of the experimental procedures.

## Author contributions

Conceptualization: I.A.T.-G., S.A., C.P.-C. and W.Z. Methodology: I.A.T.-G. for cranial windows, S.A. for GRIN lens implants; I.A.T.-G., S.A. for mini2P assembly and image acquisition; I.A.T.-G. for analysis of calcium imaging data; S.A. for analysis of behavior with D.L.C.; I.A.T.-G for analysis of synchronized behavior and neuronal activity data; I.A.T.-G., S.A., C.P.-C. and W.Z. wrote the manuscript.

## SUPPLEMENTARY FIGURES

**Supplementary Figure 1.**
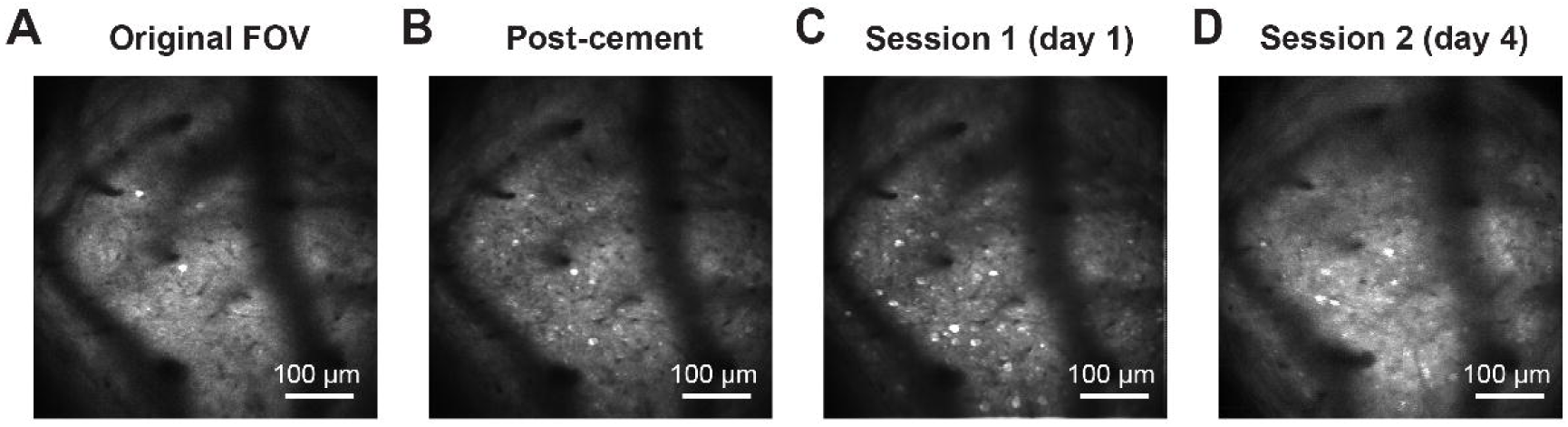
Recovery of the selected field of view across imaging sessions. **(A)** Representative Mini2P imaging FOV selected during mounting of Mini2P scope. **(B)** Same FOV after baseplate cementing **(C, D)** Same FOV recovered during two subsequent imaging sessions three days apart. Note that corresponding anatomical landmarks remain recognizable across the four recordings, providing a qualitative example of FOV recovery following repeated microscope removal and reattachment.

**Supplementary Figure 2.**
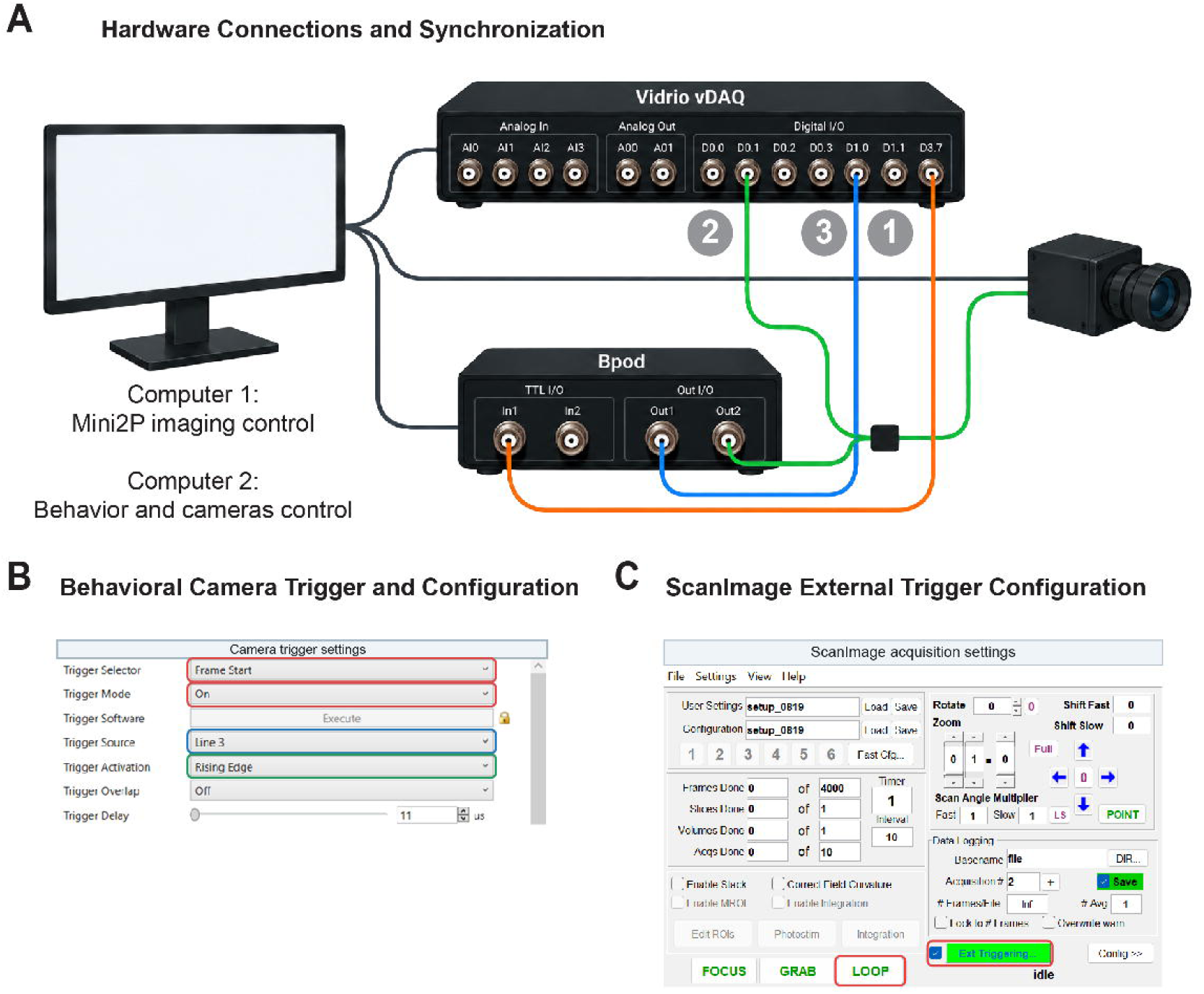
Hardware connections and trigger settings for synchronized Mini2P and behavioral acquisition. **(A)** Schematic of the computer interfaces, Vidrio vDAQ, Bpod behavioral controller, and externally triggered behavioral camera. (1) Bpod OUT1 connects to vDAQ D1.0 to initiate and terminate ScanImage acquisition on rising edges. (2) vDAQ D0.1 is connected to Bpod OUT2 and the camera through a shared connection, and (3) vDAQ D3.7 is connected to Bpod IN1. The computer connections are simplified schematically; in the experimental setup, camera acquisition and Bpod task control used a separate laptop as described in the Methods. **(B)** Behavioral camera trigger configuration, showing Frame Start as the trigger event, Trigger Mode set to On, Trigger Source assigned to Line 3, and Trigger Activation set to Rising Edge. **(C)** ScanImage Basic 2023.1.0 acquisition interface, highlighting the LOOP control and enabled external triggering required to await Bpod-controlled acquisition start/stop signals.

